# Computational array reconstruction with accumap for spatial transcriptomics

**DOI:** 10.64898/2025.12.19.695576

**Authors:** Conrad Oakes, Lior Pachter

## Abstract

Several optics-free spatial transcriptomics assays that make use of diffusion of reads from beads have recently been developed, and provide a proof of principle for high-throughput, untargeted, spatial transcriptomics. These approaches leads to a challenging computational array reconstruction problem. We show that an statistical inference method, which we term accumap, outperforms the current approach which is to run UMAP on the bead count matrix. The accumap method makes use of predicted distances to reconstruct bead positions and can improve the accuracy of reconstruction by a factor of 3 in the diffusible Slide-Tag simulation framework.

## Introduction

Measurement of cell distance by sequencing using the diffusion of beads has recently been recognized as a viable approach to optics-free spatial transcriptomics (Abdulraouf et al., 2024; Hu et al., 2025; Liao et al., 2024). Unlike imaging based spatial transcriptomics methods that rely on preselected probes (and therefore are currently limited to assaying a few thousand genes (Moses and Pachter, 2022)) or bead arrays with known barcode locations (which are expensive to produce (Liu et al., 2020)), optics-free spatial transcriptomics enables, in principle, untargeted transcriptomics with high spatial resolution at low-cost. However, a key challenge for optics-free spatial transcriptomics is the computational array reconstruction problem, i.e. the reconstruction of spatial location from count-based distance information. While distance geometry tools provide numerous approaches to reconstructions of points from distances, the spatial transcriptomics diffusion-based data provide unique and substantive challenges: distances are not directly measured, but rather counts between beads that decrease in number with distance in a non-linear manner, the data are sparse and noisy, and the measurements are high-dimensional.

The computational array reconstruction problem (CAR), is as follows: given two types of signal, sender and receiver, the input consists of a rectangular matrix with counts for the number of times each sender bead is observed with each receiver bead (see Methods). Some of the currently proposed methods allow for a single bead to be both a sender and a receiver (Abdulraouf et al., 2024), while others use two distinct sets of sender and receiver beads (Hu et al., 2025). The algorithm currently used for reconstructing the points for the CAR problem by (Abdulraouf et al., 2024; Hu et al., 2025; Liao et al., 2024) is UMAP (McInnes et al., 2018). Unlike other single-cell genomics dimension reduction problems, the CAR solution two-dimensional. Thus, methods such as UMAP, which are, in principle, optimized for embedding high-dimensional point sets in 2D, may be suitable. However, we hypothesized that UMAP may not be the ideal algorithm for the CAR problem since it is known to distort distances (Chari and Pachter, 2023) and does not make any use of known physical parameters about the system, such as the diffusion rate.

To solve CAR, we therefore studied the relationship between bead distance and read counts, and then proceeded to implement an iterative optimization procedure that directly minimizes squared error. We show that our approach, which we call accumap, greatly improves recontruction accuracy, and, importantly, improves the neighborhood reconstruction of cells which is crucial for spatial transcriptomics.

## Results

We first tested accumap on data simulated using the framework of (Hu et al., 2025), in which sender and receiver beads are distinct. To better reflect the real data generated in that study, we not only relied on the (Hu et al., 2025) framework but adjusted the number of beads to 10,000 of each type and the diffusion coeffecient to 52 um based on the reported experimental full width half maximum of 123um. To determine bounds on the performance our approach, we first tested accumap using the ground truth relationship between counts to distance. In this “perfect data” setting, we found that a single cycle of accumap improved the accuracy of cell spatial reconstruction by a factor of 2 with respect to UMAP, from a mean square error of 278 to 104 (Supplemental Table 1), while also reducing the maximum squared error between two points from 5170 to 714. As spatial transcriptomic analysis relies on neighborhood analysis, we also examined the concordance between the nearest neighbors in the accumap corrected embedding to the UMAP embedding. We found accumap has an average Jacccard distance of 0.05 for the nearest 10 neighbors versus 0.19 for UMAP, where a Jaccard distance of zero is a perfect match (Supplemental Fig. 1c).

**Fig. 1.**
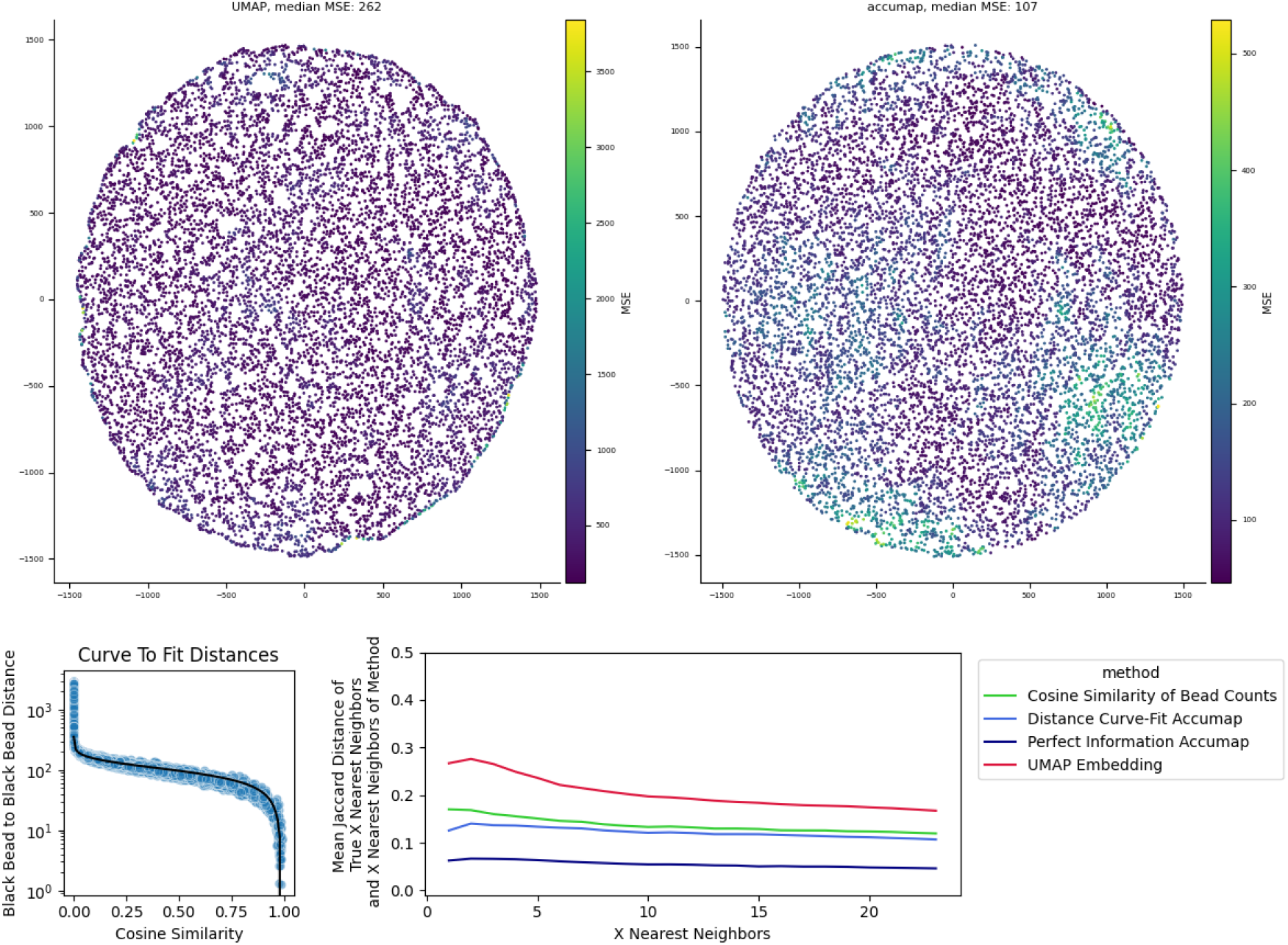
The results of accumap on a simulated dataset. A) Shows the resulting UMAP and accumap bead locations, with each bead colored by the mean squared error of its distance to every other bead. B) Shows the cosine similarity to distance curve used to fit the function that created the distance matrix for accumap. C) Average Jaccard distance between the X nearest neighbors of the true locations and the X nearest neighbors of the reconstructed array (or, in the case of cosine similarity, the X highest cosine similarities)

Encouraged by these proof-of-concept results, we next tested an approximated distance matrix generated by fitting a function relating the cosine similarity between two receiver beads and the distance between them to a small subset of 100 beads, and then applying this function to the overall 10,000 bead simulated dataset (Fig. 1b, Methods). Using this distance matrix, we found that accumap improved the accuracy of cell spatial reconstruction by a factor of 2.5 compared to UMAP, going from a mean squared error of 325 to 128 (Fig. 1a), while reducing error in 98.8 percent of beads. In the analysis of neighbors, accumap has an average Jacccard distance of.12 for the nearest 10 neighbors versus.20 for UMAP, where a Jaccard distance of zero is a perfect match (Fig. 1c).

We next sought to benchmark accumap on experimental data for which ground truth is available, but found only a single dataset available, and determined it to be of exceptionally poor quality. Of the three recently published papers, only (Hu et al., 2025) had a dataset with ground truth, and our analysis of the data uncovered major barriers to making use of it for reconstruction. Outside of the closest few neighbors (approximately 15), there is no useful spatial data, and looking at cosine similarity, we found that the Jaccard distance between actual neighbors and diffusion neighbors increases when looking at more points. This is in stark contrast to to the simulation framework for the same assay, where adding more information decreases Jaccard distance and increases accuracy (Supp Fig 3). We found that relying purely on the close local information is unreliable without further correction. In the experimental data, 56 percent of beads had a closest neighbor that was not the bead with the highest cosine similarity, and even with 10 nearest neighbors 23 percent of beads were missing 3 or more of their actual nearest neighbors among the top 10 in cosine similarity. In fact, the UMAP embedding of the real data increases the average Jaccard distance of the nearest 1 to 12 neighbors compared to assigning them purely on the basis of cosine similarity (Supp Fig 3). This calls into question its utility as a spatial dataset, as there is minimal reliable information about the neighborhood regardless of method used, even if distances can be roughly approximated by both accumap and standard UMAP. However, the fact that these trends are consistent across all replicates in their dataset suggests that the general theory behind the distance matrix core to accumap’s correction, that cosine similarity to distance correlations can be reused across datasets, is viable. Further improvements upon the technology (barcodes that better diffuse, varying ratios of sender to receiver, etc) could create a system predictable enough for accumap to be usable.

## Discussion

The accumap method greatly improves on the current standard approach to computational array reconstruction by making use of the predictable nature of diffusion between receiver beads. The general framework of the method is amenable to many different ways of determining the true distance between receiver beads, as the input for the method is a user-defined distance matrix. This could be based on fitting a function to the data, as demonstrated here, and can include distance or neighbor filters.

Our method development has also revealed that optical-free spatial transcriptomics can, in principle, provide high spatial resolution. However, widespread adoption of such methods will require better control over diffusion to allow for calibration of the diffusion count matrices.

## Methods

### Simulation

Simulations were conducted using the testing framework described in (Hu et al., 2025), as they also published ground truth experimental data on which to apply the resulting methods. To better recapitulate the eventual experimental work, we adjusted the simulation parameters to use 10,000 sender and 10,000 receiver beads (the experimental data have 12,786 and 12,030 post quality control filtering, respectively), each with a UMI of 2500 (the experimental data has an average of 2498) and a diffusion constant of 53 (calculated from the reported FHWM of 123 um). We also conducted the simulation using the simulation’s initial diffusion constant of 300 (Supp Fig 1) and with the original bead counts (Supp Fig 2).

### Fitting Distances

As the resulting single cell data only connect biological reads to receiver beads, it is not useful to directly measure a diffusion curve mapping sender to receiver beads. Instead, we fit a curve connecting the cosine similarity of the counts of every receiver bead to the distance between them. The resulting function was then used to create an expected distance matrix between each receiver bead. A minimum distance of 10um between every bead was enforced in the cases where the function outputs a physically impossible distance (i.e., negative).

### Algorithm

Accumap requires some initial embedding of distances to correct; while these could be constructed by randomly placing beads in locations, it greatly speeds up the correction process to begin with an embedding constructed by a dimensionality reduction method; here we used UMAP.

## Code and data availability

### Acquisition of experimental data and simulation framework

The experimental data and simulation framework from Hu et al. (2025) were obtained from their github repository.

### Simulation and Analysis Scripts

For details and scripts on this processing, see https://github.com/pachterlab/OP_2025_2/blob/main/analysis_scripts/New_Simulation.ipynb.

### Experimental Data Analysis Scripts

For details and scripts on this processing, see https://github.com/pachterlab/OP_2025_2/blob/main/analysis_scripts/Real_Data_Exploratory.ipynb

## Algorithm 1

accumap

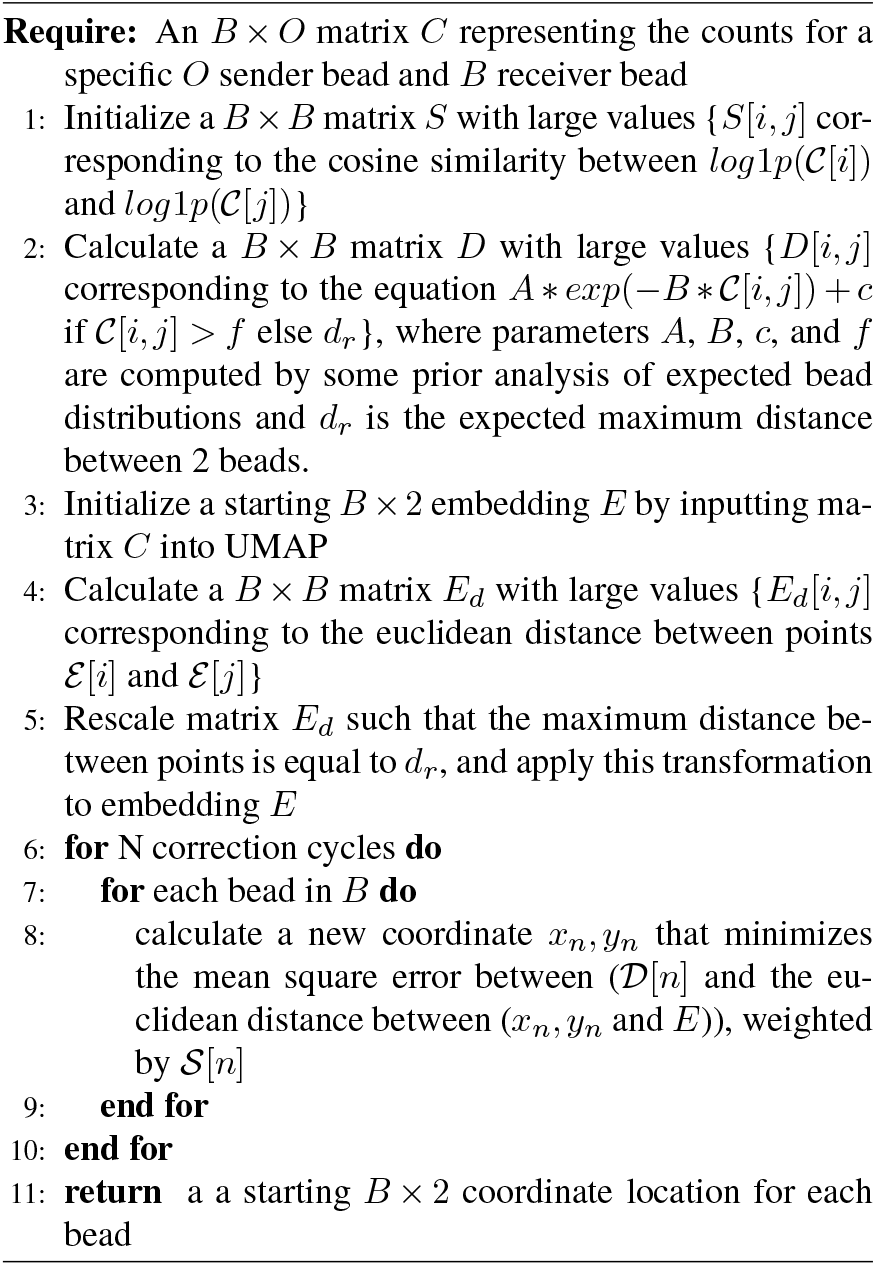

## Notes

### Competing Interest Statement

The authors have declared no competing interest.

### Summary of Updates

Updated title to include method name.

## References

Abdulraouf Abdulraouf, Weirong Jiang, Zihan Xu, Zehao Zhang, Samuel Isakov, Tanvir Raihan, Wei Zhou, and Junyue Cao. Optics-free spatial genomics for mapping mouse brain aging. bioRxiv, 2024.

Tara Chari and Lior Pachter. The specious art of single-cell genomics. PLOS Computational Biology, 19(8):e1011288, 2023.

Chenlei Hu, Mehdi Borji, Giovanni J Marrero, Vipin Kumar, Jackson A Weir, Sachin V Kammula, Evan Z Macosko, and Fei Chen. Scalable spatial transcriptomics through computational array reconstruction. Nature Biotechnology, pages 1–7, 2025.

Hanna Liao, Sanjay Kottapalli, Yuqi Huang, Matthew Chaw, Jase Gehring, Olivia Waltner, Melissa Phung-Rojas, Riza M Daza, Frederick A Matsen IV, Cole Trapnell, et al. Optics-free reconstruction of 2d images via dna barcode proximity graphs. bioRxiv, 2024.

Yang Liu, Mingyu Yang, Yanxiang Deng, Graham Su, Archibald Enninful, Cindy C. Guo, Toma Tebaldi, D. Zhang, Dongjoo Kim, Zhiliang Bai, Eileen Norris, Alisia Pan, Jiatong Li, Yang Xiao, Stephanie Halene, and Rong Fan. High-spatial-resolution multi-omics sequencing via deterministic barcoding in tissue. Cell, 2020.

Leland McInnes, John Healy, and James Melville. Umap: Uniform manifold approximation and projection for dimension reduction. arXiv preprint 1802.03426, 2018.

Lambda Moses and Lior Pachter. Museum of spatial transcriptomics. Nature Methods, 2022.

