## Supplemental File 1 for "Computational array reconstruction with accumap for spatial transcriptomics"

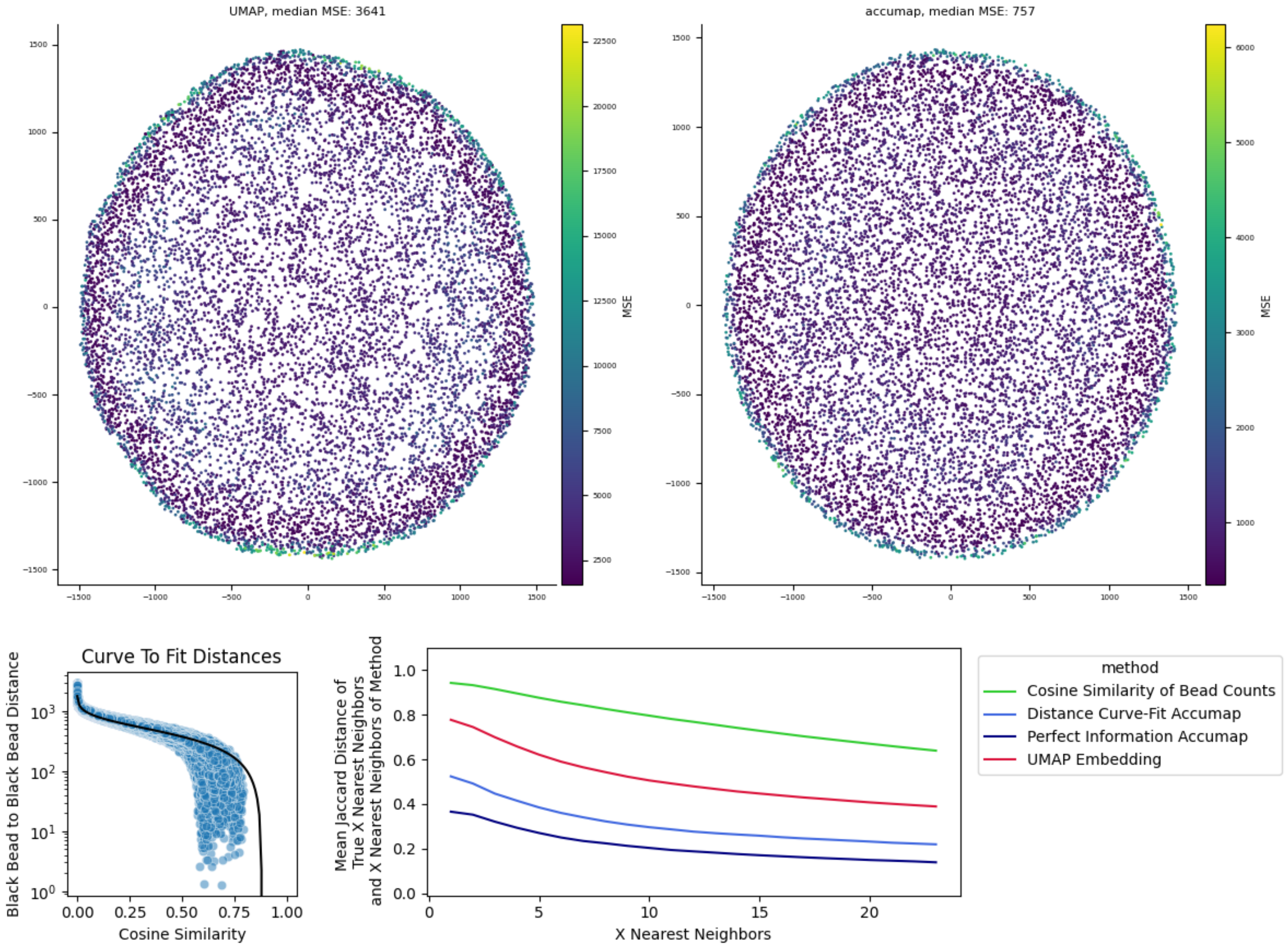


**Fig. 1.** The results of accumap on a simulated dataset using the original diffusion parameters. A) The resulting UMAP and accumap bead locations, with each bead colored by the mean squared error of its distance to every other bead. B) The cosine similarity to distance curve used to fit the function that created the distance matrix for accumap. C) Average Jaccard distance between the X nearest neighbors of the true locations and the X nearest neighbors of the reconstructed array (or, in the case of cosine similarity, the X highest cosine similarities)

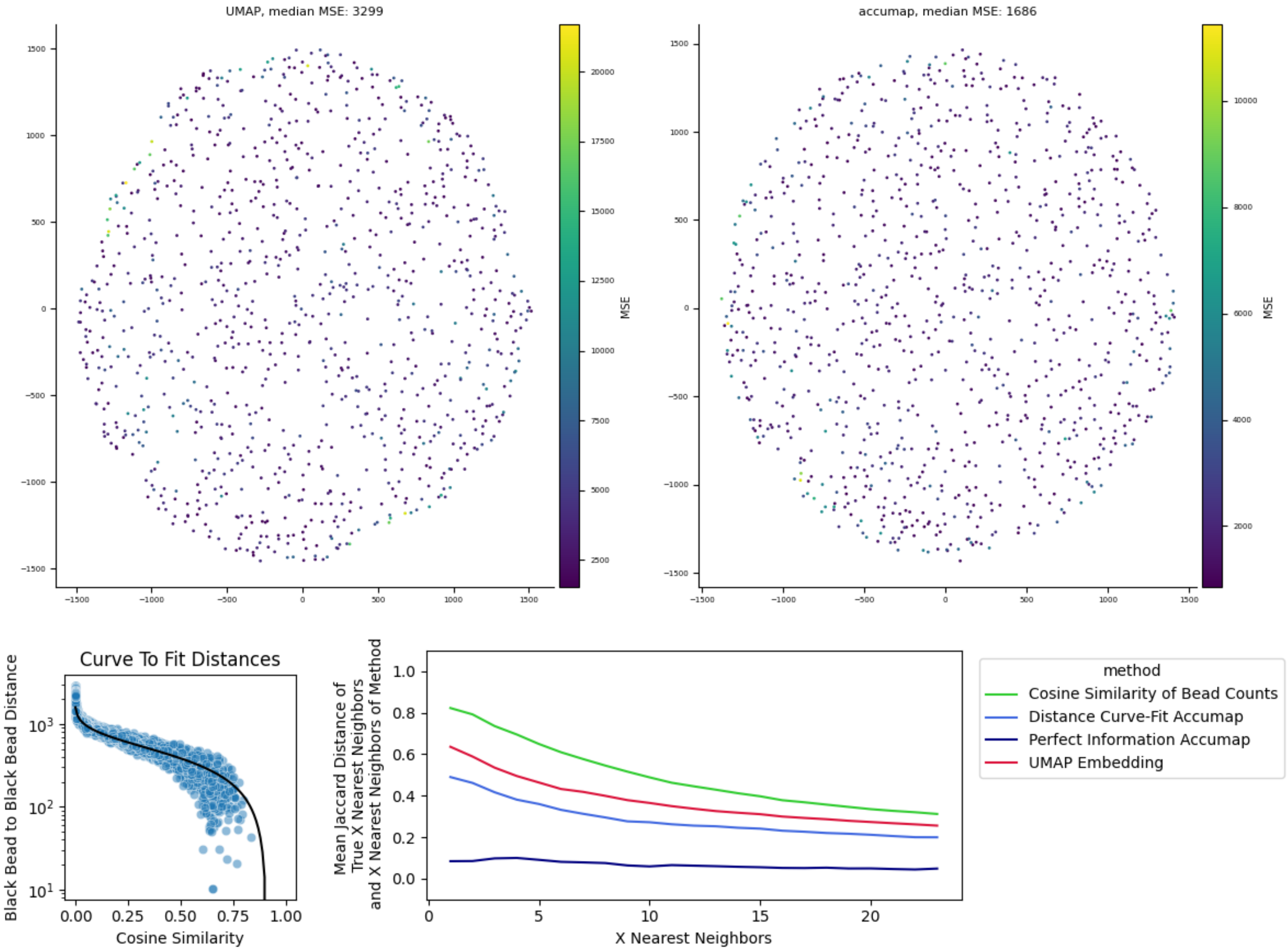


**Fig. 2.** The results of accumap on the original simulated dataset. A) The resulting UMAP and accumap bead locations, with each bead colored by the mean squared error of its distance to every other bead. B) The cosine similarity to distance curve used to fit the function that created the distance matrix for accumap. C) Average Jaccard distance between the X nearest neighbors of the true locations and the X nearest neighbors of the reconstructed array (or, in the case of cosine similarity, the X highest cosine similarities)

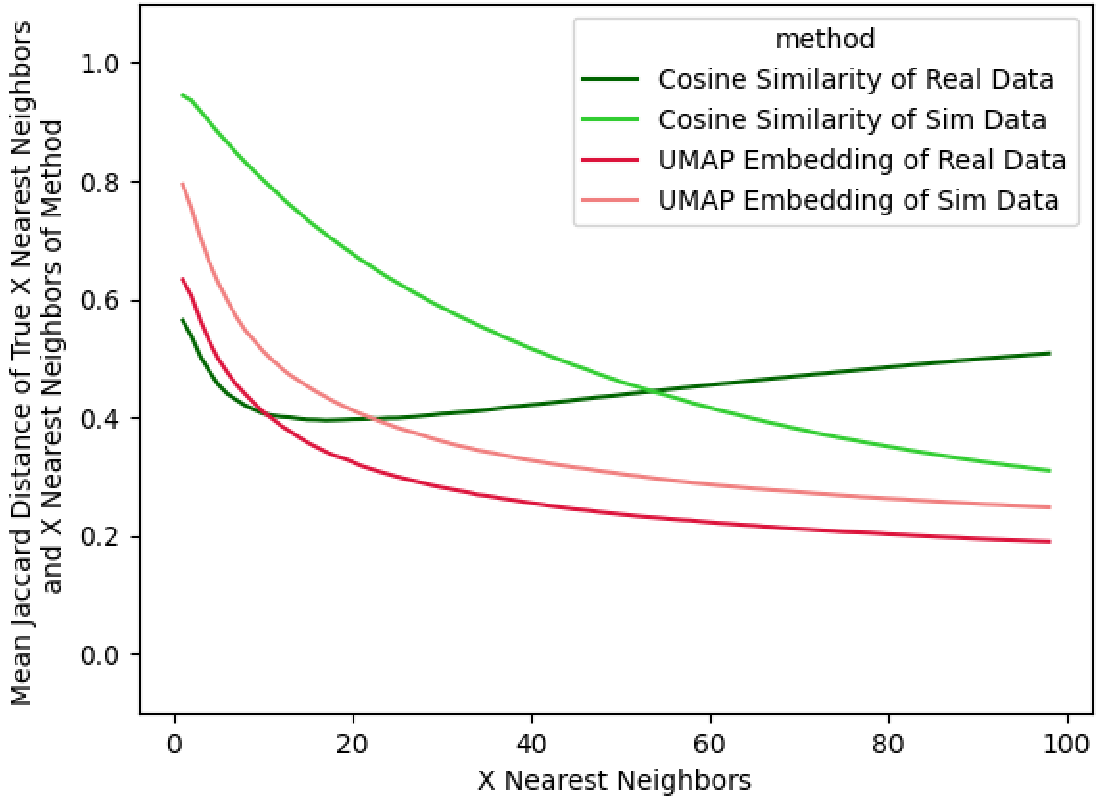


**Fig. 3.** Average Jaccard distance between the X nearest neighbors of the real or simulated dataset and the X nearest neighbors of the UMAP reconstructed array (or, in the case of cosine similarity, the X highest cosine similarities)

| Dataset | Method | Cycles | Average MSE per bead | Median MSE per bead | Average Jaccard Distance per Bead |
|---|---|---|---|---|---|
| Original Simulation | UMAP | NA | 4132 | 3300 | 0.37 |
| Original Simulation | accumap (perfect distance) | 1 | 410 | 354 | 0.07 |
| Original Simulation | accumap (fit distance) | 10 | 2069 | 1686 | 0.29 |
| New Simulation | UMAP | NA | 325 | 263 | 0.2 |
| New Simulation | accumap (perfect distance) | 1 | 167 | 145 | 0.05 |
| New Simulation | accumap (fit distance) | 10 | 126 | 108 | 0.12 |

**Table 1.** Benchmarking error metrics for accumap and UMAP in simulation.
